# Recognition of copy-back defective interfering rabies virus genomes by RIG-I triggers the antiviral response against vaccine strains

**DOI:** 10.1101/2022.03.24.485606

**Authors:** Wahiba Aouadi, Valérie Najburg, Rachel Legendre, Hugo Varet, Lauriane Kergoat, Frédéric Tangy, Florence Larrous, Anastassia V. Komarova, Hervé Bourhy

## Abstract

Rabies virus (RABV) is a lethal neurotropic virus that causes 60,000 human deaths every year around the world. A typical feature of RABV infection is the suppression of type I and III interferon (IFN)-mediated antiviral response. However, molecular mechanisms leading to RABV sensing by RIG-I-like receptors (RLR) to initiate IFN signaling remain elusive. Here, we showed that RABV RNAs are recognized by RIG-I (retinoic acid-inducible gene I) sensor resulting in an IFN response of the infected cells but that this global feature was differently modulated according to the type of RABV used. RNAs from pathogenic RABV strain, THA, were poorly detected in the cytosol by RIG-I and therefore mediated a weak antiviral response. On the opposite, we revealed a strong interferon activity triggered by the RNAs of the attenuated RABV vaccine SAD strain mediated by RIG-I. Using next-generation sequencing (NGS) combined with bioinformatics tools, we characterized two major 5’copy-back defective interfering (5’cb DI) genomes generated during SAD replication. Furthermore, we identified a specific interaction of 5’cb DI genomes and RIG-I that correlated with a high stimulation of the type I IFN signaling. This study indicates that RNAs from a wild-type RABV poorly activate the RIG-I pathway, while the presence of 5’cb DIs in vaccine SAD strain serves as an intrinsic adjuvant that strengthens its efficiency by enhancing RIG-I detection and therefore strongly stimulates the IFN response.

**Highlights:**

- RABV pathogenic strain replication *in vitro* is characterized by the absence of defective interfering genomes thus induces a weak RLR-mediated innate immunity antiviral response.
- RABV vaccine attenuated strain shows a high release of 5’ copy-back defective interfering genomes during replication *in vitro* and therefore enhances a strong antiviral response upon infection.
- RIG-I is the main sensor for RABV RNA detection within cells.

## Introduction

The innate immune response provides the first line of defense against viral infection. The cell-associated proteins known as pattern recognition receptors (PRRs) recognize the non-self motifs within viral products known as pathogen-associated molecular patterns structures (PAMPs) to trigger the release of IFN and proinflammatory antiviral cytokines.

Among PRRs, RLRs are RNA sensors localized in the cytosol. To date, this receptor family encompasses three members of helicases: RIG-I, melanoma differentiation-associated protein 5 (MDA5), and laboratory of genetics and physiology 2 (LGP2). RLRs have two common domains: i) a central DExD/H helicase domain and, ii) a carboxy-terminal domain (CTD). RIG-I and MDA5 have two amino acid terminal domains, caspase activation, and recruitment (CARDs). Upon RNA agonist binding to RIG-I or MDA5, CARDs domains interact with mitochondrial antiviral signaling protein (MAVS), mediate the downstream signal transduction and potent IFN release. IFN activates neighboring cells and via JAK/STAT pathway stimulating the expression of interferon-stimulated genes (ISGs). LGP2 lacks the CARD domain and is believed to be the fine-tuning of immune response by inhibiting RIG-I and supporting MDA5 sensing by stabilizing its interaction with RNA (Bruns et al., 2014; Sanchez David et al., 2019). Members of the RLR family are known to detect viral infections. RIG-I is the major sensor for RNA viruses belonging to *Orthomyxoviridae* (influenza A virus), *Paramyxoviridae* (Measles virus), and *Rhabdoviridae* (vesicular stomatitis virus) (Kell and Gale, 2015; Linder et al., 2021; Rehwinkel and Gack, 2020). MDA5 detects members of *Picoronaviridae* (encephalomyocarditis virus), *Coronaviridae* (SARS-CoV-2), and *Calciviridae* (murine norovirus-1) (Deddouche et al., 2014; McCartney et al., 2008; Yin et al., 2021).

Molecular characterization of RIG-I RNA ligands reveals its interaction to 5’-triphosphate dsRNA which is abrogated by the capping of the 5’-end. Cellular RNAs harboring 2’-O methyl group on the first nucleotide (N1) prevents its interaction with the RIG-I sensor which controls the immune tolerance of cellular RNAs (Hornung et al., 2006; Rehwinkel and Gack, 2020; Schuberth-Wagner et al., 2015). MDA5 recognizes the internal duplex structure and interacts preferentially to high molecular weight double-stranded RNA extracted from virus-infected cells and triggers IFN antiviral response (Pichlmair et al., 2009; Wu et al., 2013). MDA5 like RIG-I discriminates self-from virus-derived dsRNA lacking 2’-O methylation (Züst et al., 2011). Structural characterization of RIG-I and MDA5/ligands interaction indicates filament formation using protein-protein contacts to stack along dsRNA in head to a tail arrangement (Luo et al., 2011; Wu et al., 2013).

There are many types of viral RNA structures listed as being detected by RLR (Rehwinkel and Gack, 2020). Among them, the 5’cb DI genomes are probably the strongest inducers of the innate immunity response due to their double-stranded stem-loop-like structures harboring a 5’-triphosphate extremity (Baum and García-Sastre, 2011; Linder et al., 2021; Mura et al., 2017; Sanchez David et al., 2016; Strahle et al., 2006). The precise mechanism of production of these truncated forms of viral genomes by the viral polymerase is not fully understood. Whatever, they are released when the viral polymerase detaches from the template (breakpoint position: BP) and attaches to the nascent strand (reinitiation site: RI), which is then copied back producing 3’ RNA terminus with perfect complementarity to its 5’ end (Lazzarini et al., 1981). RNA viruses have evolved synergistic strategies to counteract the host’s innate immune response to sustain infection. This is the case of RABV, one of the most dangerous neurotropic zoonotic viruses causing acute encephalitis in mammals in developing countries and resulting in 60,000 human deaths every year around the world. RABV belongs to the *Rhabdoviridae* family, *Lyssavirus* genus, and possesses a negative-sense single-stranded RNA genome encoding five proteins: nucleoprotein (N), phosphoprotein (P), matrix (M), glycoprotein (G), and polymerase (L). N, P and M proteins are major antagonists of the host type-I IFN pathway and enable viral evasion of the innate antiviral response. The N protein plays an important role in the evasion of IFN response in the brain by counteracting the activation of RIG-I sensor (Masatani et al., 2010, 2013). The P protein inhibits the expression of IFN genes and ISGs by i) blocking the phosphorylation of interferon regulatory factors (IRF) 3 and IRF7; ii) preventing the IFN-I stimulated JAK/STAT pathway by the retention of activated STATs in the cytoplasm; and iii) antagonizing cytokine activated STAT3-STAT1 heterodimers (Brzózka et al., 2006; Harrison et al., 2020; Rieder et al., 2011; Wiltzer et al., 2012). In addition, RABV M protein can target RelAP43, a member of the NF-κB family, thus inducing inhibition of NF-κB signaling and reduction of expression of IFN-β genes (Besson et al., 2017; Ben Khalifa et al., 2016; Luco et al., 2012). Further, M protein cooperates with P protein to modulate the JAK-STAT pathway (Sonthonnax et al., 2019). However, few studies have so far shed light on the RLR recognition of RABV RNAs upon infection. Indeed, *in vitro* studies using RABV genetically engineered expressing little P (RABVΔP) efficiently induces IFN release (Hornung et al., 2006). In addition, RNAs isolated from RABVΔP did induce IFN expression in RIG-I overexpressing cells and this effect was strongly abrogated by the RIG-I dominant-negative mutant. Moreover, the dephosphorylation of viral RNA suppresses IFN induction, thus suggesting that RABV 5’-pppRNAs are specifically sensed by RIG-I and thus triggered IFN response (Hornung et al., 2006).

In the present study, we compared RNA molecular patterns of RLR pathway modulation induced by RABV field isolate and a vaccine strain. We used the previously validated RLR affinity purification approach combined with NGS (Chazal et al., 2018; Sanchez David et al., 2016) and identified the RNA virus-ligand signature on either RIG-I or MDA5 proteins during RABV replication. We demonstrated that IFN response induced by RLR RABV RNA recognition was principally mediated by RIG-I. 5’cb DI viral genomes that enhance RIG-I detection and therefore strongly stimulate the IFN response were exclusively produced by the RABV vaccine strain.

## Results

### RNAs purified from RABV-infected cells are immunoactive

To determine whether RABV RNAs induce IFN-mediated antiviral response, we used the previously validated ISRE reporter cell line which is human embryonic kidney-293 (HEK293) cells stably expressing firefly luciferase under the control of a promoter sequence containing five IFN-Stimulated Response Elements (ISRE) (Lucas-Hourani et al., 2013). Total viral RNAs were extracted from infected human neuroblastoma cells (SK.N.SH) with either the virulent cell culture-adapted canine RABV field isolate, THA or the SAD vaccine strain and then transfected into the ISRE reporter cell line. As expected, ISRE expression was induced when the reporter cells were transfected with 5’-pppRNAs (5’3P), low molecular weight (LMW) poly(I:C), and high molecular weight (HMW) poly(I:C). Interestingly, RNA molecules of THA and SAD induced statistically significant (p <0.01, one-way ANOVA using Tukey method) activation of ISRE promoter expression (Figure 1A). In addition, SAD RNA molecules extracted from infected cells showed a statistically significant stronger activation (p=0.007, one-way ANOVA using Tukey method) of ISRE expression than observed for parent strain THA.

**Figure 1:**
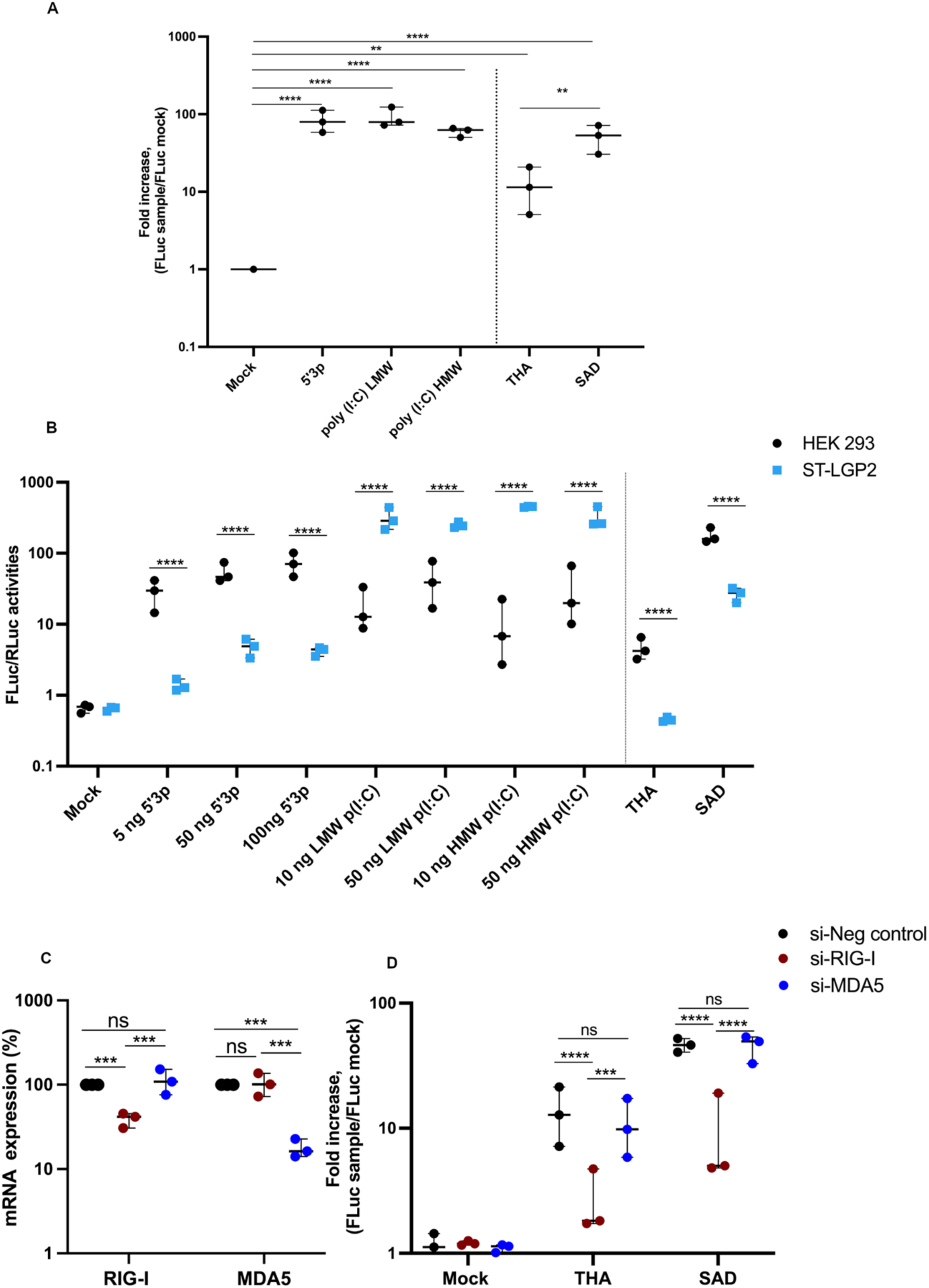
RNAs purified from RABV infected cells are immunoactive: (**A**)SK.N.SH cells were infected with THA or SAD at MOI of 0.5 for 72h, viral RNAs were extracted, transfected into ISRE-reporter cells, and firefly luciferase expression was monitored. (**B**)IFN-β promoter expression in HEK293 cells (black) or in HEK293 expressing one-Strep tagged-LGP2 ST-LGP2 (blue) transfected with 50 ng of THA, SAD RNAs or with each of the indicated synthetic RLR ligands. (**C**) Relative expression of *RIG-I* and *MDA5* mRNAs in STING37 silenced with non-targeting si-RNAs si-Negative control (black), targeting RIG-I (si-RIG-I in red) or MDA5 (si-MDA5 in blue). (**D**) The indicated silenced ISRE-reporter cells were transfected for 24h with 50 ng of total RNAs extracted form THA -or SAD-infected cells followed by luciferase assay measurement. The results are represented as fold increase of ISRE expression compared to Mock non transfected cells. The experiments were performed three times and represented as median with 95% confidence interval. P values were calculated using one way ANOVA with Tukey’s multiple comparisons. Non-significant (n.s.) were indicated. *p<0.05, **p< 0.01, ***p<0.001, ****P<0.0001

### RIG-I is the main sensor of RABV infection in HEK293 cells

LGP2 is the third member of the RLR family and it is known to inhibit RIG-I and amplify MDA-5-dependent responses, respectively (Bruns et al., 2014; Sanchez David et al., 2019). Therefore, to further evaluate the implication of RIG-I and/or MDA5 in RABV RNA detection and IFN signaling, we used HEK293 cell line stably over-expressing One-STrEP-tagged LGP2 (ST-LGP2) (Sanchez David et al., 2016). We have previously demonstrated that LGP2 overexpression enhances MDA5- and blocks RIG-I-specific RNA ligands activation of IFN-β promoter signaling (Sanchez David et al., 2019). We evaluated IFN-β expression in either ST-LGP2 or parental HEK293 cells transfected with commonly used synthetic RNAs or with RABV RNA extracted from THA-or SAD-infected cells together with reporter plasmid expressing firefly luciferase (Fluc) under the control of IFN-β promoter. As expected, we observed that the transfection of ST-LGP2 with 5’3P synthetic RNA significantly less activated the expression of Fluc compared to the parental HEK293 cells, whereas the transfection of ST-LGP2 with MDA5-specific ligands (LMW or HMW) increased the activity of the IFN-β promoter. Similarly, to 5’3P synthetic RNA, THA and SAD RNA molecules transfected in HEK293 induced a strong increase of IFN-β promoter activity, whereas ST-LGP2 transfected cells showed suppression of IFN signaling compared to those observed in parental HEK293 cells (Figure 1B) indicating that RIG-I is mainly implicated in RABV RNA detection upon infection.

To explore in more detail the differential involvement of RLR (RIG-I or MDA5) in RABV RNA sensing, the ISRE reporter cell line was treated with non-targeting siRNAs (si-Neg control), targeting RIG-I (si-RIG-I) or MDA5 (si-MDA5). Transient silencing of RIG-I and MDA5 significantly (p ≤0.001, one-way ANOVA using Tukey method) reduced the level of mRNA for RIG-I and MDA5 by ∼61% and ∼82%, respectively as compared to si-Negative control cells (Figure 1C). When the same cells were transfected with RABV RNAs (THA or SAD), only RIG-I silenced reporter cells showed strongly impaired ISRE promoter activation (p <0.0001, one-way ANOVA using Tukey method), while MDA5 silencing did not affect the signaling (Figure 1D).

Altogether, these results demonstrate that RIG-I is the main cytosolic PRR that detects RABV RNAs to mediate IFN signaling.

### During THA and SAD infections immunoactive RNA ligands bind to RIG-I and modulate IFN response

To evaluate the IFN stimulation activity of RLR-bound ligands from RABV-infected cells, we took advantage of previously validated HEK293 cells (ST-RLRs) expressing STrEP-tagged RIG-I (ST-RIG-I) and MDA5 (ST-MDA5) (Sanchez David et al., 2016).

First, we tested whether RLR overexpression would influence RABV infection. ST-RIG-I and ST-MDA5 cells were infected at different multiplicities of infections (MOIs) either with the pathogenic RABV THA strain or the SAD vaccine strain and compared to the negative control ST-CH cell line expressing STrEP-tagged Red Fluorescent Protein Cherry (Sanchez David et al., 2016). We first ascertained the efficacy of RABV replication in ST-RLR cells. To this aim, the virus growth was monitored at 16, 24, and 48h post-infection by quantification of genomic RNA by qPCR (**Figure S1**). THA replicated less efficiently than SAD in ST-RLR cells and especially at a low MOI of 0.1 UFF/cell. In addition, SAD replication was efficient even at the early stage of replication. Furthermore, THA and SAD replications were not altered in cells expressing additional copies of RIG-I or MDA5 compared to ST-CH negative control cells. Then we evaluated the activity of RNA molecules bound to RLR. To this aim, ST-RIG-I, ST-MDA5, and ST-CH cell lines were infected by either THA or SAD at MOI of 0.5 and harvested at 24h post-infection. The efficiency of the purification of cherry, RIG-I, and MDA5 proteins obtained from total cell lysates (input) and after affinity purification (output) was controlled by western blotting (Figure 2A).

**Figure 2:**
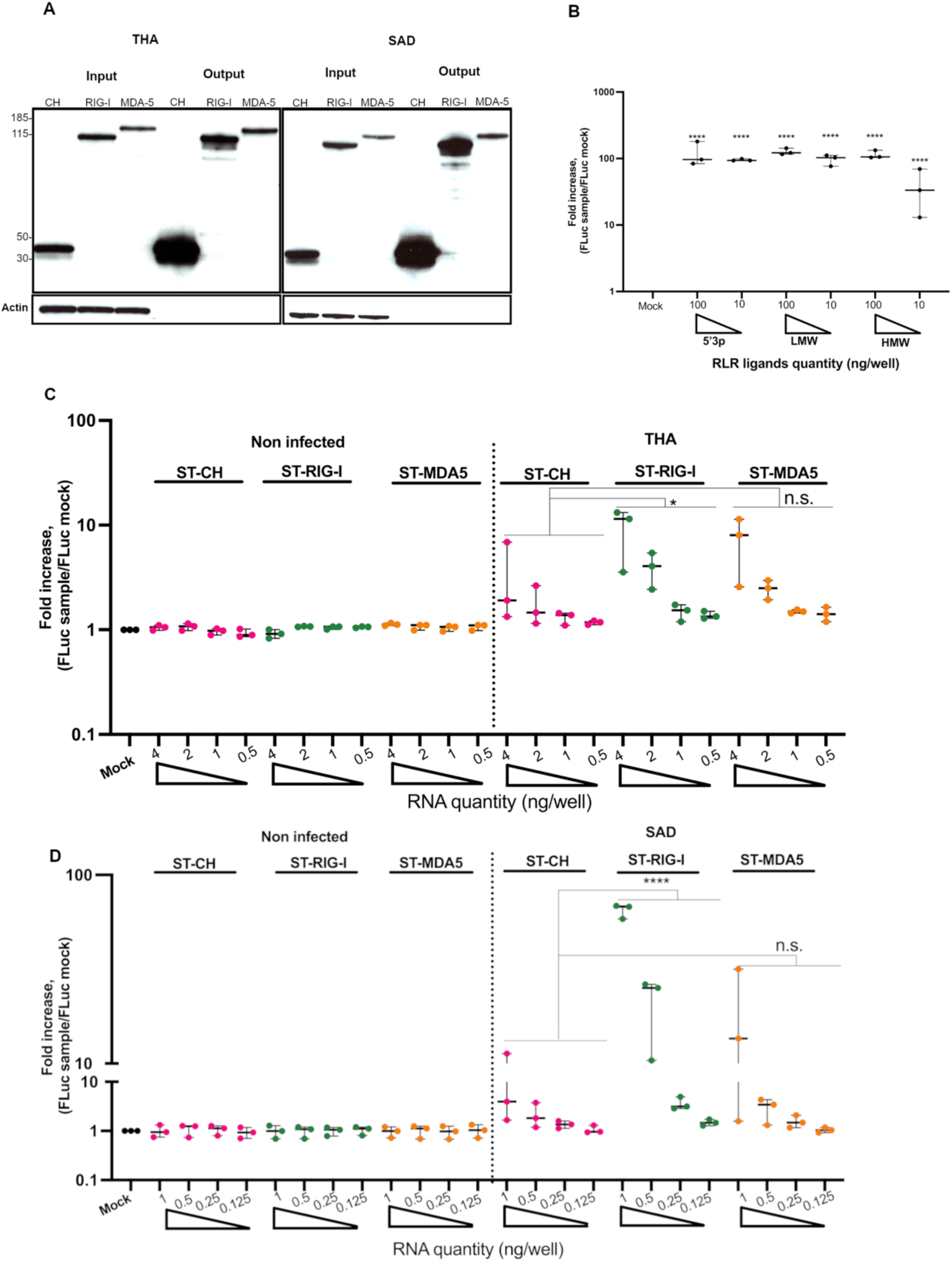
THA and SAD ligands bound to RIG-I are immunoactive: (**A**)western blot analysis of RLR protein expression in ST-RLR infected cells. Total cell extracts (input) were affinity-purified using Strep-tagged beads (output). The western blot was performed using α-STrEP-Tag (upper blot) or α-ß-actin (bottom blot) antibodies. Luciferase expression of STING37 cells transfected with a quantity gradient of RLR synthetic ligands (**B**), THA (**C**)-or SAD-RNAs (**D**) co-purified with strep-tagged negative control cherry (CH represented in pink), RIG-I (green), or MDA5 (orange) extracted from ST-RLR cells-non-infected or infected at MOI of 0.5 for 24h. Experiments were done in triplicate and represented as median with 95% confidence interval. The results are represented as a fold increase of ISRE expression compared to Mock non-transfected cells. P values were calculated using one-way ANOVA with Tukey’s multiple comparisons. Non-significant (n.s.) were indicated. *p<0.05, **p< 0.01, ***p<0.001, ****P<0.0001

Then, RNAs co-purified with RIG-I, MDA5, or CH were extracted and transfected into the ISRE reporter cell line to assess the activation of type-I IFN signaling. First, the ISRE promoter activation was first controlled by transfecting synthetic RNAs. As expected, 5’3P, HMW poly(I:C), and LMW poly(I:C) largely stimulated ISRE expression (Figure 2B). These data validated our transfection approach. RIG-I-specific RNAs extracted from THA-infected cells induced a light but significant (p=0.029, single-step method) stimulation of ISRE expression as compared to the negative control CH-specific RNAs (Figure 2C). RIG-I-specific SAD RNA induced a strong and significant (p=0.00007, single-step method) ISRE promoter activity in a dose-dependent manner as compared to negative control CH-specific RNAs (Figure 2D). For both viral strains, MDA5-specific RNAs showed no significant stimulation of the ISRE promoter activity. These data suggest that RNA extracted from THA and SAD infected cells present molecular patterns that are absent in non-infected cells and that are preferentially recognized by RIG-I.

### The molecular pattern of RABV recognition by RIG-I and MDA5

To further characterize RLR bound RABV RNAs, ST-RLRs cells were infected with THA or SAD at MOI of 0.5. The cells were harvested after 24h, and RLRs bound RNAs were purified using affinity chromatography. Total RNAs from input cell extract, as well as RLRs-bound RNA output, were extracted. Input and output RNAs were subjected to NGS followed by bioinformatics analysis using the previously described protocol (Chazal et al., 2018). NGS provided high output in the order of 12 to 90 million total reads per sample with around 0.3 and 0.29% mapped to THA and SAD genomes, respectively (Table S1). The fold change of RLR binding was obtained by normalizing: i) mean abundance of reads (coverage) of either RIG-I or MDA5-ligands by mean read coverage of corresponding input samples, ii) nonspecific RNAs binding were discarded by normalizing RLRs read coverage to reads obtained with cherry negative control. NGS data analysis revealed no visible enrichment of read-abundance in RLR association of negative- and positive-sense viral RNAs upon THA infection in our experimental conditions (Figure 3A). Interestingly, the NGS study of the RABV vaccine strain, SAD, showed a significant coverage and much higher enrichment on MDA5 of negative-sense RNA along the whole viral genome than with positive-sense viral RNA. Thus, MDA5 may be engaged in recognition of the SAD negative-sense genome. Conversely, studies analyzing RNA bound to MDA5 in measles virus-infected cells (also negative-sense single-stranded RNA virus) reporting that MDA5 interacts with only positive-strand viral RNA (Runge et al., 2014; Sanchez David et al., 2016). However, despite viral RNA/MDA5 interaction, we failed to observe any statistically significant stimulation of ISRE promoter in reporter cells in our experimental conditions (Figure 2D).

**Figure 3:**
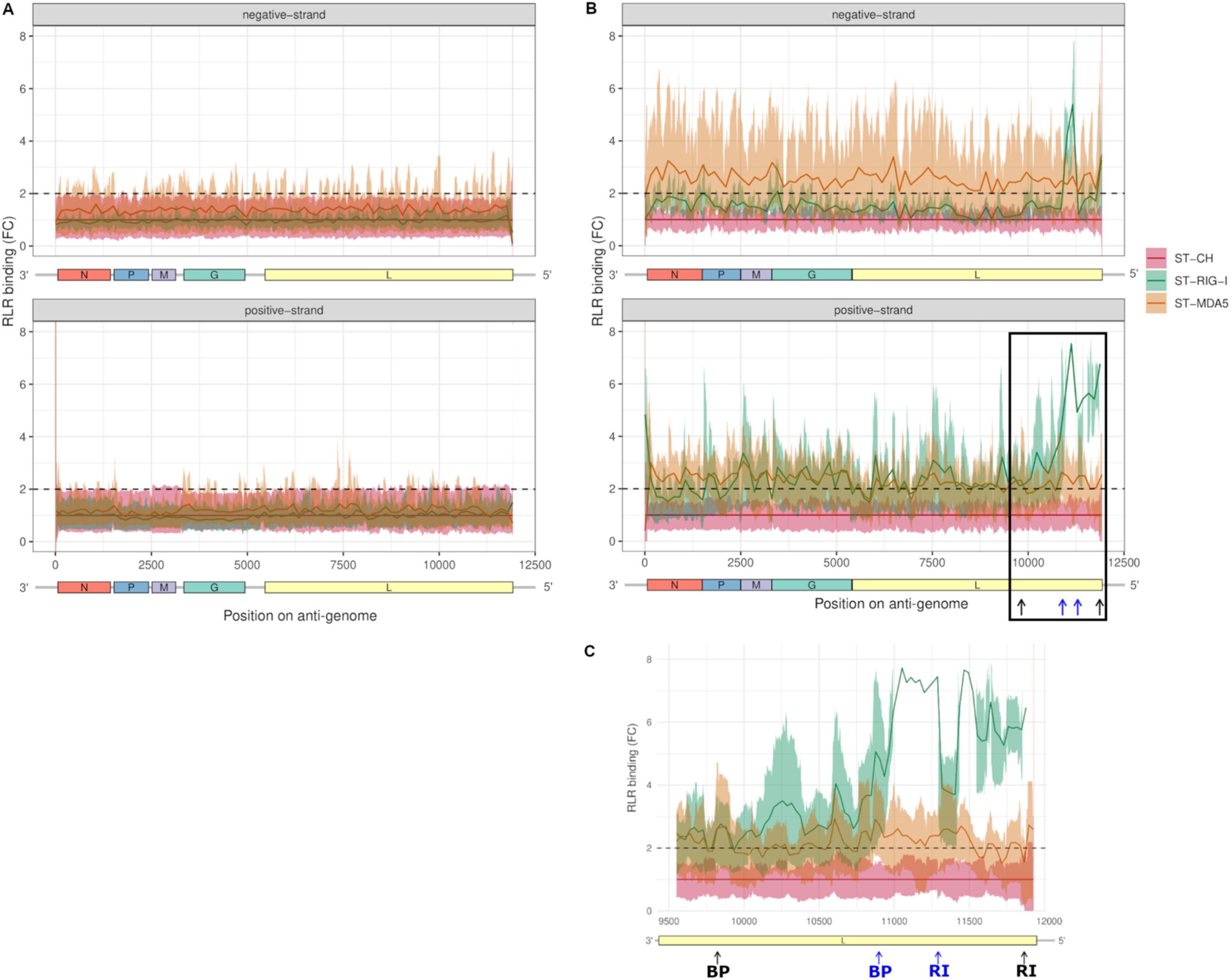
Molecular pattern of RABV recognition by RIG-I and MDA5: RLR cells were infected with THA or SAD at MOI of 0.5 for 24h. (**A**) THA or (**B**) SAD total RNAs extracts (Input) or co-purified (output) with ST-CH (pink), ST-RIG-I (green), or ST-MDA5 (orange) are subjected to strand-specific NGS analysis. The read sequences obtained were mapped to THA (**A**) or SAD (**B**) genome references in negative orientation (upper panel) or positive orientation (bottom panel) and are depicted on the y-axis as RLR binding fold change (FC) at each specific position along genome sequence represented on the x-axis. Schematic annotations of THA (**A**) or SAD genome (**B**) underline the x-axis. (**C**) Zoom enlargement of the black-framed panel in B. Three independent experiments are represented as smooth curves. Black and blue arrows depicted the breakpoints, BP, and reinitiation sites, RI, for 5’ cb DI-2170 and 5’cb DI-1668, respectively. The fold change of RLR binding is obtained by i) normalizing the read coverage of output RNAs co-purified with RIG-I, MDA5 or CH by their input extract, ii) normalized read coverage of output samples were divided by the mean of the coverage of negative control CH, at each genomic position.

Conversely, the analysis of RIG-I-specific RNA ligands purified from SAD-infected cells revealed a significant enrichment of viral negative- and positive-sense RNAs (Figure 3B) and in particular the 5’ and 3’ extremities of genomic and antigenomic RNAs, respectively. These results are in concordance with the RLR silencing experiment suggesting that RIG-I is the main actor in RABV RNAs recognition (Figure 1) and with the statistically significant stimulation of ISRE promotor by the SAD RNAs bound to RIG-I (Figure 2D). Thus, NGS data analysis of RLR-specific RNAs upon RABV infection suggested that the extremities of the genomic and antigenomic viral RNAs would play a particular role in the immune stimulation induced by SAD vaccine strain.

To further explore the RNA primary sequence motifs of RLR ligands that could explain SAD RNA recognition by RIG-I and or MDA5, we analyzed the AU content of RIG-I and MDA5 specific reads as described previously (Sanchez David et al., 2016). We observed that MDA5 binding to SAD negative-strand genomic RNAs correlated with high AU content (> AU content of SAD genome 0.55) (p<0.001, Cohen’s d=0.53) (Figure S4D). However, RIG-I SAD negative-strand ligands were characterized by a lower AU-rich content sequence than that observed for MDA5 ligands suggesting that RIG-I binding did not correlate with RNA primary sequence (Figure S4B). For the positive-strand SAD RNA species, we did not detect any preference of binding of AU-rich regions by RIG-I or MDA5 (Figure S4A and C). In conclusion, our results suggest preferential binding of MDA5 to AU-rich regions of viral RNA but RIG-I ligand interaction to AU-rich sequence was not observed suggesting that RIG-I more presumably recognizes specific RNA secondary or tertiary structures.

### Characterization of RIG-I protein specific SAD 5’cb DI viral genomes

We questioned if RIG-I specific negative- and positive-sense viral RNA observed during SAD infection (Figure 3B) could be attributed to 5’ cb DI viral genomes. Indeed, 5’ cb DI viral genomes are known strong RIG-I ligands (Baum et al., 2010; Mura et al., 2017; Sanchez David et al., 2016). We applied DI-tector algorithm to search for the presence of 5’cb DI viral genomes in THA and SAD NGS data sets (Beauclair et al., 2018). We failed to detect any 5’cb DI genome in ST-RLR cells infected with THA. Interestingly, using the same approach, we identified two 2170- and 1668-nucleotides long 5’ cb DI viral genomes in SAD-infected cells Table 1). 5’cb DI-1668 exhibited statistically significant (p =0.04, one-way ANOVA) read abundance in the RIG-I output samples as compared to the negative control ST-CH. We failed to detect any statistically significant read enrichment for 5’cb DI-2170 in ST-RIG-I output samples comparing to the negative control ST-CH cells (Table 1).

**Table 1:** DI-tector results of NGS data analysis of SAD-infected cells.

| 5' cb DI name | Samples | Length (nt) | BP position | RI position | ST-CH |  |  | ST-RIG-I |  |  | P values | ST-MDA5 |  |  | P values |
| --- | --- | --- | --- | --- | --- | --- | --- | --- | --- | --- | --- | --- | --- | --- | --- |
|  |  |  |  |  | R1 | R2 | R3 | R1 | R2 | R3 |  | R1 | R2 | R3 |  |
| **DI-2170** | **Input** | 2170 | 9823 | 11865 | 4 | 0 | 4 | 7 | 0 | 0 | n.s. | 0 | 0 | 4 | n.s. |
| **DI-1668** |  | 1668 | 11292 | 10898 | 2 | 0 | 0 | 0 | 3 | 0 | n.s. | 4 | 2 | 7 | n.s. |
| **DI-2170** | **Output** | 2170 | 9823 | 11865 | 0 | 0 | 0 | 14 | 2 | 0 | n.s. | 0 | 0 | 0 | n.s. |
| **DI-1668** |  | 1668 | 11292 | 10898 | 0 | 0 | 0 | 73 | 11 | 10 | * | 6 | 0 | 0 | n.s. |

Data sets were generated from RNA samples obtained from SAD-infected ST-Cherry (ST-CH), ST-RIG-I or ST-MDA5 cells before (Input) or after purification (Output).
For each 5'cb DI, the breakpoint position (BP) and reinitiation site: (RI) are identified.
P values were calculated using one-way ANOVA with Tukey's multiple comparisons between counts of reads obtained from ST-RIG-I or ST-MDA5 vs negative control (ST-CH) from three independent replicates (R1; R2; and R3). Non-significant (n.s.) were indicated. \*p<0.05, \*\*p<0.01, \*\*\*p<0.001, \*\*\*\*p<0.0001

We further studied molecular organization of the detected 5’cb DI-2170 and 5’cb DI-1668 DI viral genomes and juxtaposed them on the Figure 3B with SAD RIG-I/RNA NGS data representation. 5’cb DI-2170 resulted from a breakpoint (BP) at position 9823 of the full-length viral genome and reinitiation (RI) at position 11865. 5’cb DI-1668 generated from a BP position at 11292 and RI at position 10898 (Table 1, Figure 3C). Complete DI-genome sequences are indicated in table S3. We observed that both negative- and positive-sense 5’cb DI viral genomes were enriched on RIG-I in SAD-infected cells (Figure 3B). Indeed, similar to the full-length viral genome, 5’cb DI genomes are replicated *via* the production of anti-genomes which are used as templates. 5’ cb DI viral genomes are characterized by the presence of trailer- and its reverse complementary-sequence at 5’ and 3’ extremities which hybridize forming a stem-loop structure (Figure S3) that has been shown to stimulate RIG-I (Mura et al., 2017; Schlee et al., 2009).

Our data demonstrate for the first time the lack of detectable amount of cb DI viral genomes in the RABV field isolate (THA) which may fine-tune the outcome of viral infection and persistence. However, the SAD vaccine strain presents an important source of 5’cb immunogenic RNAs recognized *via* the RIG-I.

### Validation of 5’cb DI viral genomes production by RABV SAD strain using a conventional approach

To further validate the presence of 5’cb DI viral genomes in SAD-infected ST-RLR cells, we applied a universal RT-PCR analysis on total RNA extracted from ST-RLR cells infected with SAD at MOI of 0.5 for 24h. Using universal 5’cb DI genome-specific primers as described previously (Mura et al., 2017; Pfaller et al., 2014; Shingai et al., 2007), two DNAs fragments were detected on agarose gel of approximately 0.7 Kb and 1.2 Kb corresponding to 5’cb DI-1668 and 5’cb DI-2170, respectively (Figure 4A). Furthermore, Sanger sequencing of the two amplicons confirmed the characteristics of the 5’ cb DI genomes i.e., BP and RI sites (Table S3). To further extend our findings, the presence of 5’cb DI genomes were examined in human neuroblastoma cells (SK.N.SH), a more relevant cell model to study RABV infection and pathogenesis. These cells were infected with either RABV field isolate THA or RABV vaccine strain SAD. Using universal primers, only SAD infected cells demonstrated the presence of two PCR fragments corresponding to DI-1668 and DI-2170 (Figure 4B). Furthermore, we failed to detect any 5’cb DI genomes from THA infected cells using optimized THA-specific primers (Figure 4C). Thus RT-PCR analysis of 5’ cb DI viral genomes in SK.N.SH cells corroborated with DI-tector study performed on our NGS data (Figure 3). Therefore, these experiments validated the presence of identical 5’cb DI viral genomes produced independently of the cell type in SAD-infected ST-RLR and human neuroblastoma cells.

**Figure 4:**
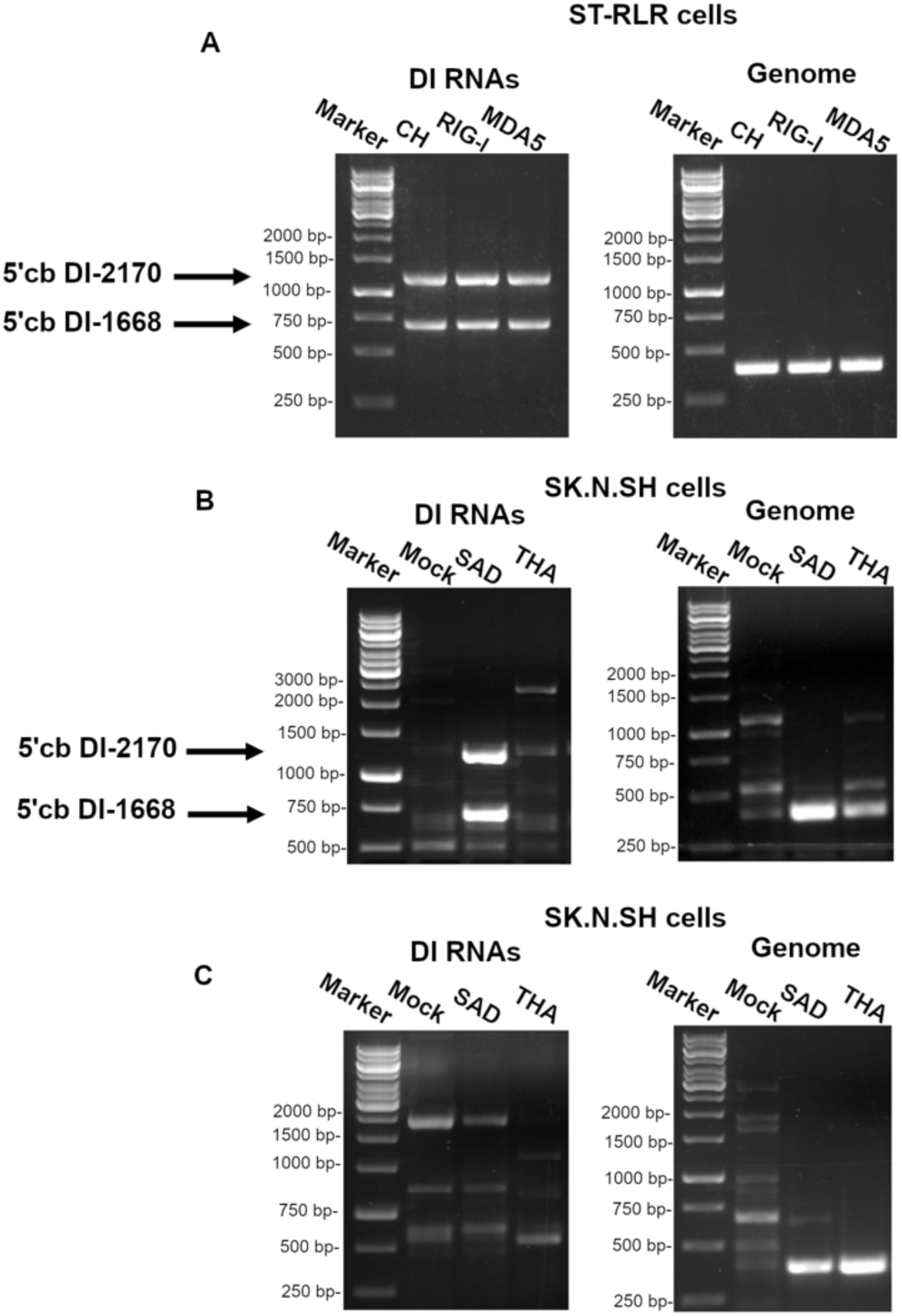
Characterization of SAD 5’ cb DI viral genomes identified by DI-tector and their validation by RT-PCR: A. Detection of 5’cb DI-1668, 5’cb DI-2170 (left panel), and SAD genome (right panel) by RT-PCR on total RNAs extracted from SAD infected ST-CH, ST-RIG-I, and ST-MDA5 cells (MOI of 0.5, 24h post-infection). RT-PCR was performed using universal 5’ cb primers (1 and 2) and full-length genome primers (2 and 3) (Table S3). B. Detection of 5’cb DI-1668, 5’cb DI-2170 (left panel), SAD and THA genomes (right panel) by RT-PCR on total RNAs extracted from infected neuroblastoma cells (SK.N.SH) at MOI of 0.5 for 24h using universal 5’ cb primers (1 and 2) and full-length genome primers (2 and 3) (Table S3). C. RT-PCR on 2 µg of total RNAs extracted from SAD or THA infected neuroblastoma cells (SK.N.SH) at MOI of 0.5 for 24h using THA-optimized 5’ cb DI primers (4 and 5) (left panel) and THA-optimized full-length genome primers (3 and 4) (right panel) (Table S3).

## Discussion

RLRs cytosolic sensors are the first line of defense that triggers the innate immune response to infection by detecting invasion of viral RNAs in the cytoplasm. Therefore, understanding the RLR signaling could help to develop antiviral therapeutics that control viral infection. Further, it could shed light on some of the mechanisms explaining the attenuation of some viruses. Several studies have performed characterization of RNA partners bound to RLRs within infected cells using various riboproteomic approaches. RNAs bound to RLRs were isolated by Co-IP or tagged-protein affinity purification and then characterized by NGS. Using these approaches, first RIG-I-specific RNA partners for negative-sense RNA viruses (Sendai, influenza, VSV, and Measles viruses) and positive-sense RNA viruses (Dengue, Zika viruses, Chikungunya) were identified (Baum et al., 2010; Chazal et al., 2018; Linder et al., 2021; Sanchez David et al., 2016).

RABV is thought to counteract IFN induction and inflammation by many ways (Faul et al., 2009; Lafon, 2008). Indeed, mice infected with attenuated RABV induced stronger inflammatory reactions than mice infected with the wild-type RABV (Wang et al., 2005). However, only a few studies on RABV RNA recognition have been performed and most of them carried in the absence of productive virus infection by transfecting cells with *in vitro* transcribed RNA or RNAs coming from RABV infected cells (Hornung et al., 2006). Therefore, we lack a deeper investigation on real RNAs signatures recognized by RLRs during RABV infection. Here, we addressed this question in the presence of active infection with two RABV strains: cell culture-adapted canine RABV field isolate from Thailand (THA) and a vaccine strain (SAD) used largely to produce live-attenuated and inactivated vaccines.

We demonstrated that RNAs molecules isolated from THA-or SAD-infected cells induced ISRE and IFN-β expression upon their transfection. But we observed that the induction of ISRE was not comparable for the two viruses, the vaccine strain induced a stronger interferon response than the field isolate did. Further, silencing of either RIG-I or MDA5 suggested RIG-I to be the major PRR for THA or SAD viral strains. These observations are in agreement with previous work indicating that RIG-I is required for the initiation of an IFN response upon cell infection with SAD L16 (Hornung et al., 2006). In addition, the LGP2 overexpression inhibited the IFN-β expression in ST-LGP2 cells transfected with RABV RNAs that confirmed the role of RIG-I in RNAs detection upon RABV infection. Our results corroborate previous studies on RIG-I implication in the detection of negative-sense RNA viruses (Sendai virus, Measles virus and Influenza virus) (Kell and Gale, 2015; Mura et al., 2017; Rehwinkel et al., 2010; Sanchez David et al., 2016) and more specifically on the previously described study performed on Vero cells transfected with plasmids encoding either for the full-length or a truncated (dominant-negative mutant, RIG-IC) RIG-I and infected with SAD-L16 strain, a SAD related strain (Hornung et al., 2006).

Using RLR/RNA pull-down approach coupled to NGS, we detected an enrichment of 5’ cb DI genomes of both negative- and positive-sense in our RIG-I-specific RNA samples and only in SAD-infected cells. Moreover, we found the evidence for the interaction of the full-length viral genome with MDA5 in SAD-infected cells. However, we failed to detect any immunostimulation activity of MDA5-specific RNA partners in type-I IFN reporter assays. *In silico* analysis showed that the genomic sequence of SAD bound to MDA5 presented an AU-rich composition. It has been reported that the AU-rich RNA species bound to MDA5 are poorer activators of ATP hydrolysis of MDA5 *in vitro* forming a stable MDA5 filament structure (Runge et al., 2014). This feature may explain the lack of immunostimulatory activity of the MDA5-SAD ligand due to their AU-rich composition reported in our study. However, this observation may be cell-specific as RABV was previously shown to induce IFN in dendritic cells of RIG-I knockout mice suggesting that MDA5 could be in that particular case involved in RABV sensing upon infection (Faul et al., 2010).

Identical pull-down/ NGS approach applied to the pathogenic field isolate THA failed to detect any viral RNA ligands specifically bound to RIG-I. The immunostimulation activity of the RNAs collected from the RIG-I pull-down sample in THA infected cells should not be considered as a discrepancy between our data. Indeed, it could be explained either by the too low sensitivity of the RLR/NGS approach when used in THA infection conditions and/or by the presence of immune active Pol3-produced endogenous RNA (Vabret *et al*., yet unpublished data), or other small self-RNAs produced by the action of the antiviral endoribonuclease RNase L on cellular RNAs. The RNase L by-products are sensed by RIG-I and therefore induce IFN-β expression (Malathi et al., 2007).

In line with our observations, DI RNA genomes were isolated from a broad range of other negative-sense RNA viruses including VSV, Sendai virus, Measles virus, influenza A virus (recently reviewed in (Ziegler and Botten, 2020). Further, DI particles have also been found in human natural infections and correlated with the course of the disease (Sun et al., 2015). Indeed, genomic analysis of influenza A virus (IAV) H1N1 isolated from a cohort of highly severe/fatal outcomes identified few DIs while isolates from a mild case of disease produced a high level of DIs (Vasilijevic et al., 2017). Moreover, the kinetics of DI accumulation and duration can predict the clinical outcome of RSV A infection in humans (Felt et al., 2021). Thus, this suggests that DIs could be considered as a new virulence marker for viral pathogenicity (Vasilijevic et al., 2017). In the case of RABV, DI particles were also observed *in vivo* in newborn mice brains inoculated intracerebrally with the highly attenuated RABV HEP Flury strain or VSV (Holland and Villarreal, 1975).

In summary, using the riboproteomic approach coupled to the NGS we provide here a significant positive correlation between the presence of 5’ cb DIs and the strong RIG-I mediated interferon response and the weak virulence of vaccine strain SAD in living cells (Figure 5).

**Figure 5:**
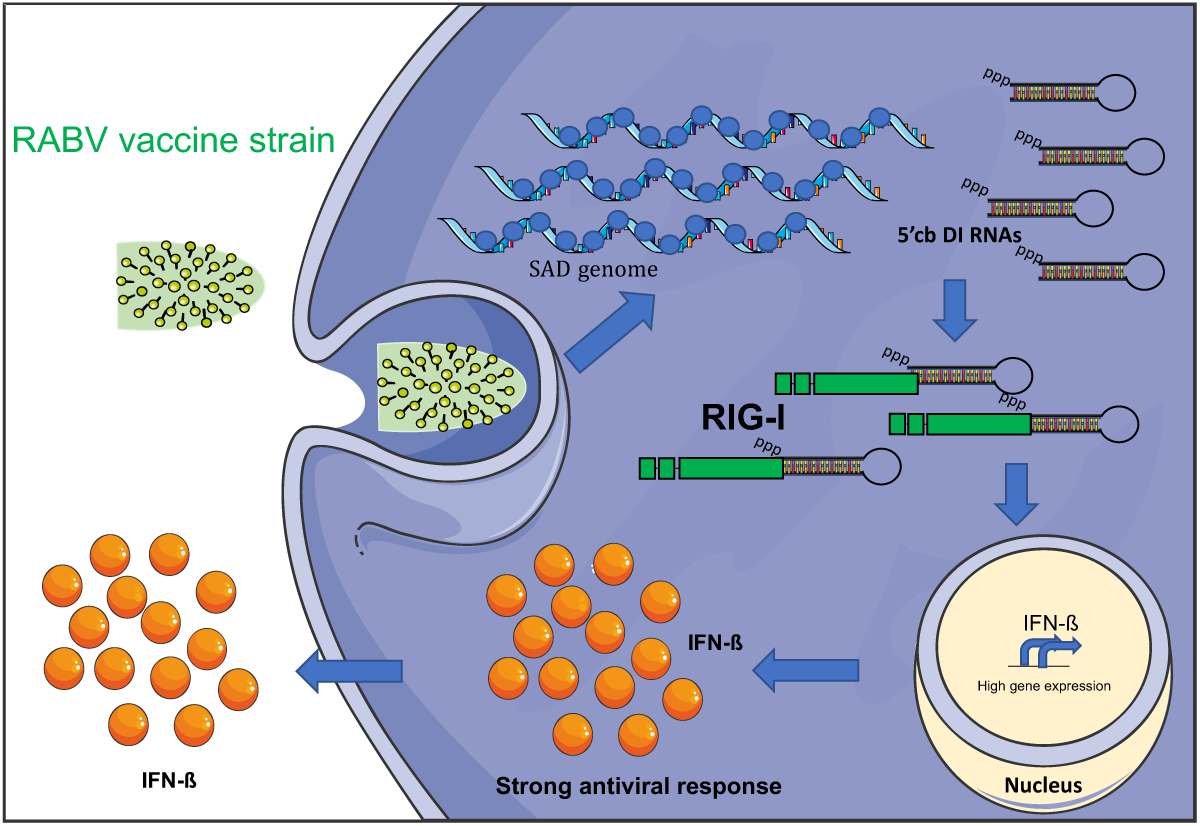
Model of SAD RNAs recognition by RIG-I and modulation of IFN response: During vaccine strain cell infection SAD genome and 5’cb DI RNAs are replicated in cytoplasm. 5’cb DI RNAs are then detected by RIG-I sensor triggering a signaling cascade that activate IFN response.

## Material and methods

### Virus and cells

SK-N-SH (human neuroblastoma, ATCC-HTB-11), HEK293 (human embryonic kidney, ATCC CRL-1573) were maintained in DMEM supplemented with 10% calf serum. ISRE-reporter HEK293 (STING37) and HEK293 expressing one strep-tag RLR cells (Lucas-Hourani et al., 2013; Sanchez David et al., 2016): ISRE reporter STIN37 and HEK293 ST-RLR cells were maintained in DMEM supplemented with 10% calf serum containing G418 at 500 mg/ml (#G8168, SIGMA, St. Louis, Missouri).

Rabies virulent cell culture-adapted canine RABV field isolate THA, (EVAg collection Ref-SKU: 014V-03194, ENA accession n° GCA_927797755) was obtained by reverse genetics using RNA extracted from 8743THA strain isolated from a man bitten by a dog in Thailand, (EVAg collection, Ref-SKU: 014V-02106, GenBank accession n° EU293121.1).

Rabies vaccine SAD B19 Bern-C strain (EVAg collection Ref-014V-01929, ENA accession n° GCA_927797725).

### RNAi experiments

STING 37 cells were plated for 24h before transfection at 37°C (2×10^5^ cells/well), non-targeting small interfering RNAs: **i**) non-targeting pool (si-control, #D-001810-10-05, smart pool: 5’UGGUUUACAUGUCGACUAA3’, 5’UGGUUUACAUGUUGUGUGA3’, 5’UGGUUUACAUGUUUUCUGA3’, 5’UGGUUUACAUGUUUUCCUA3’), **ii**) si-RIG-I-(#L-012411-00-0005, smart pool: 5’GCACAGAAGUGUAUAUUGG3’, 5’CCACAACACUAGUAAACAA3’, 5’CGGAUUAGCGACAAAUUUA3’, 5’UCGAUGAGAUUGAGCAAGA3’) or **iii**) si-MDA-5 (#L-013041-00-0005, smart pool: 5’GAAUAACCCAUCACUAAUA3’,5’GCACGAGGAAUAAUCUUUA3’, 5’UGACACAAUUCGAAUGAUA3’, 5’CAAUGAGGCCCUACAAAUU3’) were added at 40 nM in the presence of DharmaFECT 1 transfection reagent (#T-2001-03, Horizon discovery), then incubated at 37°C for 24h.

### Reverse transcription-PCR analysis

Total RNA was extracted using a RNeasy minikit (Qiagen). cDNA was generated from 2 *µ*g of total RNA by using Superscript IV vilo (#11766050, Invitrogen) and primer 2 (Table S2) in a total volume of 20 *µ*l. A total of 2 *µ*l of the resultant cDNA was then amplified with primers 2 and 3 for genome amplification and primers 1 and 2 for DI-RNA amplification (Table S2), using Phusion high-fidelity DNA polymerase (Thermo Fisher Scientific) in a total volume of 50 *µ*l (95°C for 2 min; 40 cycles of 95°C for 30 s, 55°C for 30 s, and 72°C for 1 min; and 72°C for 10 min). The products were analyzed in a 1.5% agarose gel, with a GeneRuler 1kb DNA ladder (#SM0312; Thermo Fisher scientific).

### Western blot and antibodies

Protein extracts were resolved by SDS-polyacrylamide gel electrophoresis on 4-12 % gradient NUPAGE-PAGE gel (Invitrogen) with MOPS running buffer and transferred to cellulose membranes (GE Healthcare) with the Criterion Blotter system (Biorad). The following antibodies were used: anti-STrEP-Tag (#34850, Qiagen), monoclonal anti-β-actin antibody (A5441, Sigma), HRP-coupled anti-mouse (NA9310V, GE Healthcare) or anti-rabbit (RPN4301, GE Healthcare) were used as secondary antibodies. Peroxidase activity was visualized with an ECL Plus Western Blotting Detection System (#RPN2132, GE Healthcare).

### Immunostimulation assay on STING37 and HEK 293 cells

Total virus-RNAs were extracted from SK.N.SH infected cells at MOI 0.5 using RNeasy minikit (Qiagen, #74104). ISRE reporter assay was performed in STING37 cells plated in 24 wells plate (2×10^5^ cells/well) and incubated for 24h at 37°C. The RNAs were transfected using lipofectamine 2000 reagent (#11668019, Invitrogen,) in optiMEM medium. After 24h, the transfected cells were harvested and lysed using 200 *µl* of passive lysis buffer (#E1941, Promega). Luciferase (Fluc) expression was assessed using Bright-Glo Luciferase Assay System (#E2638, Promega).

IFN-β reporter assay was assessed by transient transfection of HEK293 and ST-LGP2 cells using reporter plasmid expressing luciferase under the control of IFN-β promoter. The cells were plated (2×10^5^ cells/well) and incubated at 37°C for 24 h before transfection. The cells were transfected with 700 ng, 70 ng, and 50 ng of IFN-β, Renilla reporter plasmids, and total viral RNAs, respectively. 5’3P RNA was synthetized *in vitro* using Xba1-linearized pCINeo plasmid. T7 transcription was carried out using T7 RiboMax Express large scale RNA production kit (# P1320, Promega).

### Purification of ST-RLR/RNAs by affinity chromatography

ST-RLRs cells (15×10^6^cells) were infected at MOI of 0.5 by either THA or SAD. The cells were harvested after 24h, washed twice with cold PBS, and lysed in 2 ml lysis buffer (20 mM MOPS pH 7.4, 120 mM KCl, 0.5% igepal, 2 mM MgCl_2_, 2 mM β-mercaptoethanol) supplemented with protease inhibitor cocktail (#11873580001, Roche) and 200 unit/ml RNAsin (#N2515, Promega). The cell lysate was left on ice for 20 min with gentle mixing and then clarified by centrifugation at 13,000 rpm for 15 minutes at 4°C. For total RNA (input) extraction, an aliquot of 100 µl of each cell lysate was used in the presence of TRI reagent LS (#T3934, SIGMA). The remaining lysate supernatant was incubated for 2 h on a spinning wheel with 200 µl of high-Performance strepTactin Sepharose beads (#28-9355-99, GE healthcare). The beads were collected by centrifugation at 1600 g at 4°C for 5 min, washed three times on spinning wheel for 5 min with 5 ml washing buffer (20 mM MOPS pH 7.4, 120 mM KCl, 2 mM MgCl_2_, 2 mM β-mercaptoethanol) supplemented with protease inhibitor cocktail and 200 unit/ml RNasein. RNAs bound to RLR (output) were eluted using biotin elution buffer (IBA), extracted with TRI reagent LS and resuspended in 40 µl ultrapure RNase free water. The quality of input and output RNAs was checked by Bioanalyzer RNA pico- and nano-kits, respectively (Agilent, Santa Clara, California).

### Library preparation

RNA molecules extracted from input and output preparations were used for library preparation using the TruSeq stranded total RNA library prep kit (Illumina, San Diego, California), according to the manufacturer’s protocol. The first step of poly-A RNA isolation was omitted to analyze all RNA species present. First, the RNAs were randomly fragmented followed by random primed reverse transcription. dsDNA fragments were generated by second-strand DNA synthesis. To add specific, Illumina adapters, an adenine was added to the 3’ extremity followed by adapter ligation. The quality of all libraries was checked with the DNA-1000 kit (Agilent) on a 2100 Bioanalyzer and quantification was performed with Quant-It assays on a Qubit 1.0 fluorometer (Invitrogen). Sequencing was performed with the Illumina NextSeq500 system. Runs were carried out over 75 cycles, including seven indexing cycles, to obtain 75-bp single-end reads. Sequencing data were processed with Illumina Pipeline software (Casava version 1.9).

### Read mapping and statistical analysis

Reads were cleaned of adapter sequences and low-quality sequences using cutadapt version 3.2. Only sequences at least 25 nt in length were considered for further analysis. Bowtie version 2.1.0 (with –very-sensitive mode) was used for alignment on the reference genomes THA and SAD, respectively. Coverage was obtained using BEDTools with genomecov -d parameters (Quinlan and Hall, 2010). Analyses were performed with R version 4.0.3, and bioconductor packages ggplot2 and dplyr as described previously (Chazal et al., 2018). Read coverage of output samples were normalized by mean read coverage of their input extracts. To obtain RLR binding, the normalized output samples were normalized by the mean of the negative control (cherry samples triplicates), at each genomic position. The RLR binding were plotted using ggplot2.

### *In silico* analysis of RLR-RNAs partners

AU content was calculated in a sliding window of 200 nucleotides with a step size of one nucleotide and was compared to the mean count within this window. We apply a T-test effect size using Cohen’s d measure to compare AU content distributions between significant and non-significant positions.

### Statistical analysis

An ANOVA model including the replicate effect as blocking factor was used across the biological conditions. Pairwise comparisons were extracted using the emmeans R package so that the p-values are adjusted for multiple testing using the Tukey method.

## Data availability

The data discussed in this publication have been deposited in NCBI’s Gene Expression Omnibus (Edgar et al., 2002) and are accessible through GEO Series accession number GSEXXX.

## Acknowledgements

We acknowledge all the members of the Lyssavirus, Epidemiology and Neuropathology unit and the Vaccines-innovation laboratory at Pasteur Institute for their help and useful discussions. We thank Alexis Criscuolo (Hub Bioinformatique et Biostatistique, Institut Pasteur) for help with bioinformatics analysis.

This study was supported by ANR RAB-CAP (ANR-16-CE11-0031-01), the Institut Pasteur and the CNRS.

## Author contribution

WA performed the most of the experiments, analyzed the data and wrote the manuscript. VN performed transfection, affinity purification, and western blotting experiments. RL carried out the bioinformatics analyses. HV carried out all the statistical analysis. LK performed the library preparation. FL participated in validation experiments. HB and AVK designed the study, edited, and discussed the manuscript. All authors read and approved the final manuscript.

## Supplementary data

**Table S1:** Read count and coverage for NGS samples (related to figure 3)

| Extract | Sample | THA |  |  |  |  | SAD |  |  |  |  |
| --- | --- | --- | --- | --- | --- | --- | --- | --- | --- | --- | --- |
|  |  | Total reads | Mapped reads | % Mapped reads | Coverage |  | Total reads | Mapped reads | % Mapped reads | Coverage |  |
|  |  |  |  |  | plus | minus |  |  |  | plus | minus |
| **Output** | **ST-CH R1** | 41855395 | 99391 | 0.24 | 446 | 177 | 12556635 | 29997 | 0.24 | 114 | 744 |
|  | **ST-CH R2** | 29662451 | 49926 | 0.17 | 224 | 889 | 7374170 | 23411 | 0.32 | 897 | 573 |
|  | **ST-CH R3** | 97987753 | 235869 | 0.24 | 1049 | 431 | 15482658 | 40539 | 0.26 | 178 | 768 |
|  | **ST-MDA5 R1** | 57110195 | 140375 | 0.25 | 642 | 239 | 73953920 | 149197 | 0.2 | 684 | 252 |
|  | **ST-MDA5 R2** | 72568282 | 87252 | 0.12 | 407 | 140 | 30434181 | 129937 | 0.43 | 390 | 426 |
|  | **ST-MDA5 R3** | 37930998 | 94636 | 0.25 | 403 | 191 | 43187126 | 146331 | 0.34 | 634 | 285 |
|  | **ST-RIG-I R1** | 42748638 | 72499 | 0.17 | 331 | 124 | 69836256 | 177587 | 0.25 | 715 | 100 |
|  | **ST-RIG-I R2** | 51203231 | 53702 | 0.1 | 253 | 836 | 46901371 | 96863 | 0.21 | 421 | 167 |
|  | **ST-RIG-I R3** | 32382885 | 119162 | 0.37 | 458 | 290 | 61150678 | 155870 | 0.25 | 769 | 211 |
| **Input** | **ST-CH R1** | 65141630 | 249896 | 0.38 | 1040 | 529 | 39595348 | 124352 | 0.31 | 546 | 235 |
|  | **ST-CH R2** | 47461521 | 246756 | 0.52 | 867 | 682 | 18757555 | 43062 | 0.23 | 173 | 97 |
|  | **ST-CH R3** | 30964668 | 214868 | 0.69 | 776 | 572 | 16675786 | 52672 | 0.32 | 234 | 96 |
|  | **ST-MDA5 R1** | 32252644 | 128293 | 0.4 | 544 | 261 | 27463544 | 106863 | 0.39 | 461 | 210 |
|  | **ST-MDA5 R2** | 69251116 | 264582 | 0.38 | 992 | 670 | 30297107 | 66517 | 0.22 | 250 | 168 |
|  | **ST-MDA5 R3** | 32793430 | 136394 | 0.42 | 530 | 328 | 55207608 | 220288 | 0.4 | 981 | 403 |
|  | **ST-RIG-I R1** | 40871809 | 137001 | 0.34 | 560 | 300 | 90476392 | 280687 | 0.31 | 1 188 | 575 |
|  | **ST-RIG-I R2** | 26123818 | 66630 | 0.26 | 248 | 170 | 37798581 | 70879 | 0.19 | 274 | 171 |
|  | **ST-RIG-I R3** | 50607423 | 385424 | 0.76 | 1378 | 1042 | 29007870 | 113288 | 0.39 | 498 | 213 |

**Table S2:**
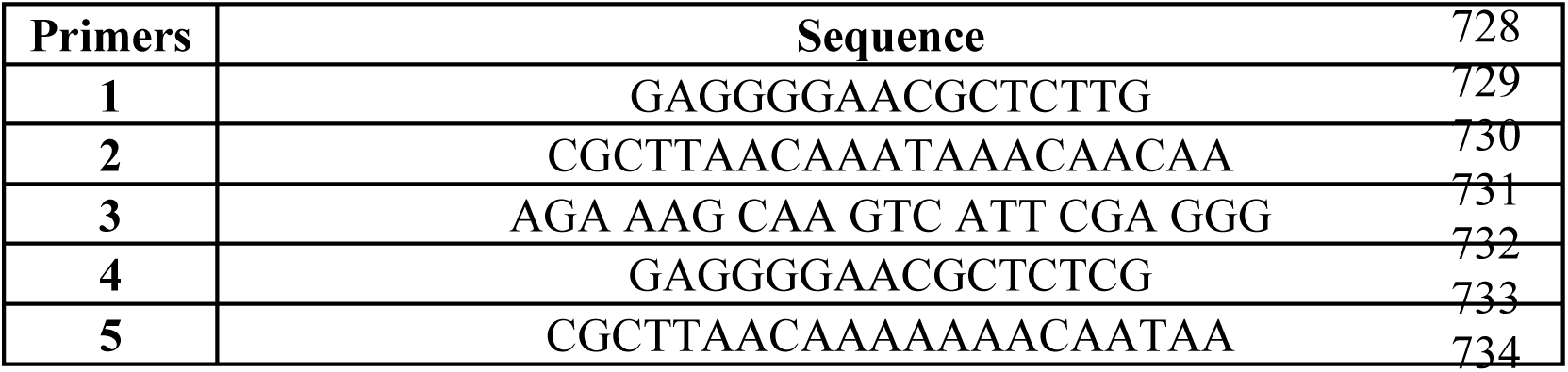
primers used in RT-PCR.

**Table S3:**
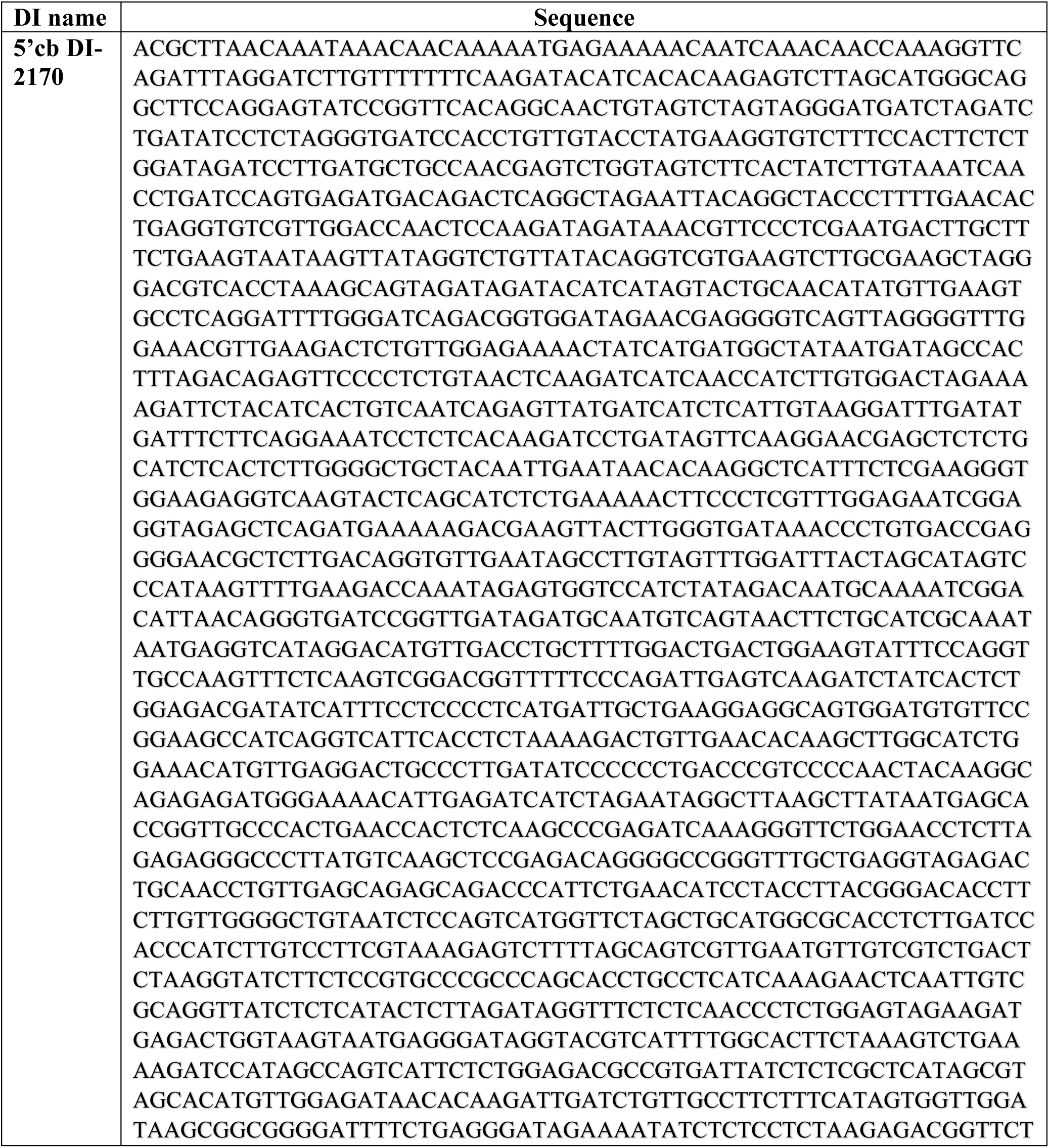
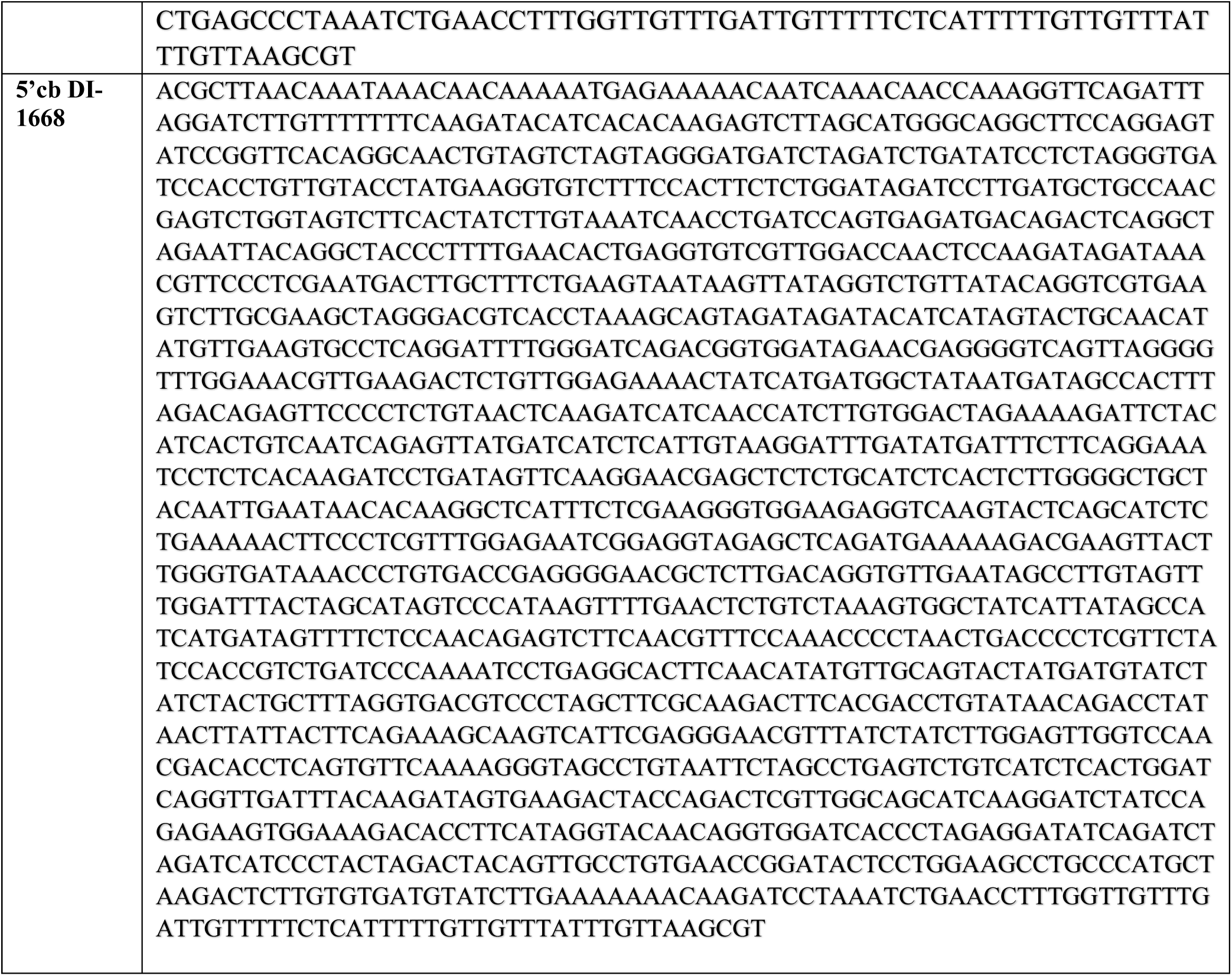
Sequences of 5’cb DI genomes.

**Figure S1:**
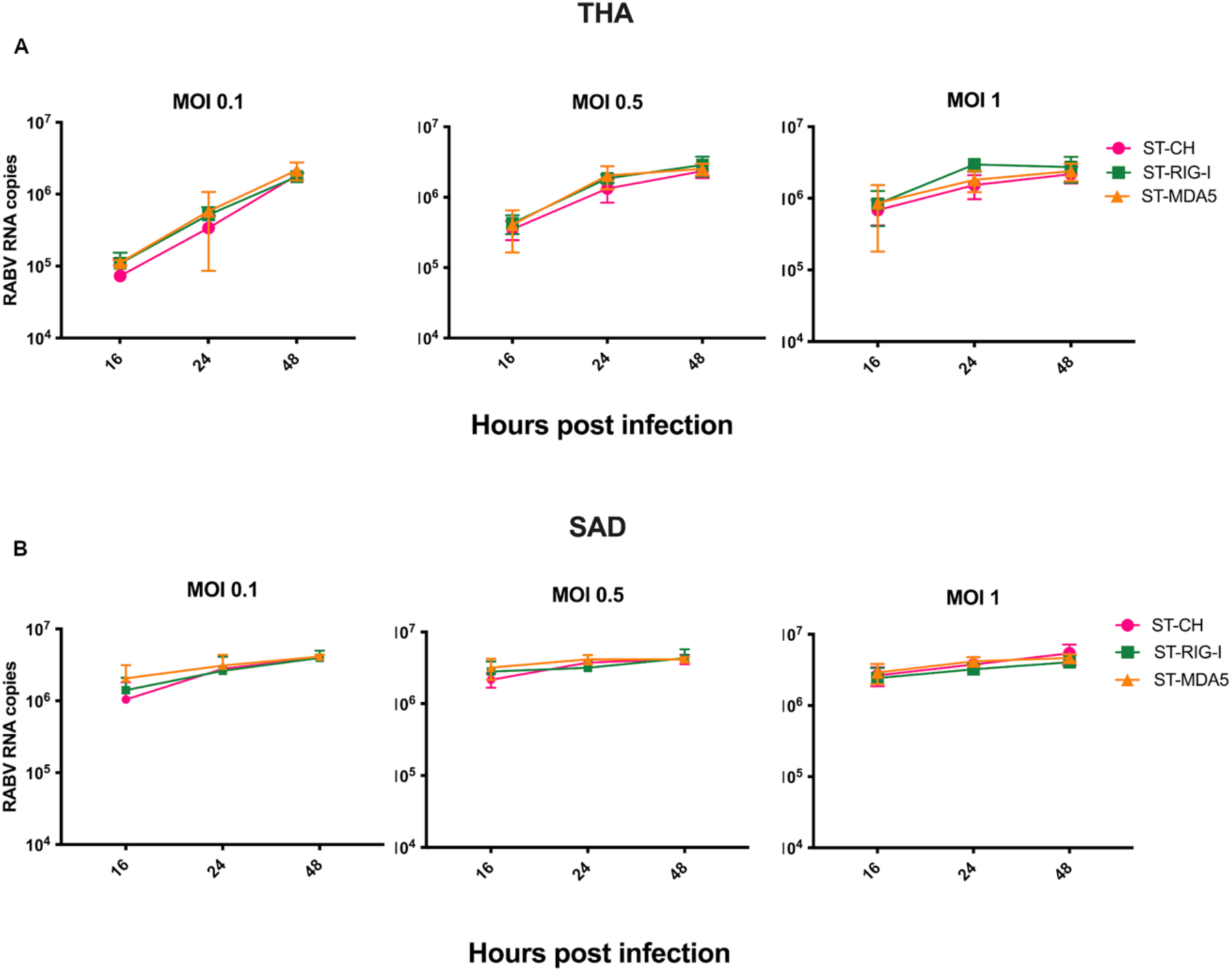
RABV replication in RLR-overexpressing cells. ST-RLR cells were infected with THA (A) or with SAD (B) at different MOIs (0.1, 0.5. or 1). The growth kinetics was monitored 16, 24, and 48h post-infection by qPCR of RABV genomic RNA at position from 1130 to 1247.

**Figure S2:**
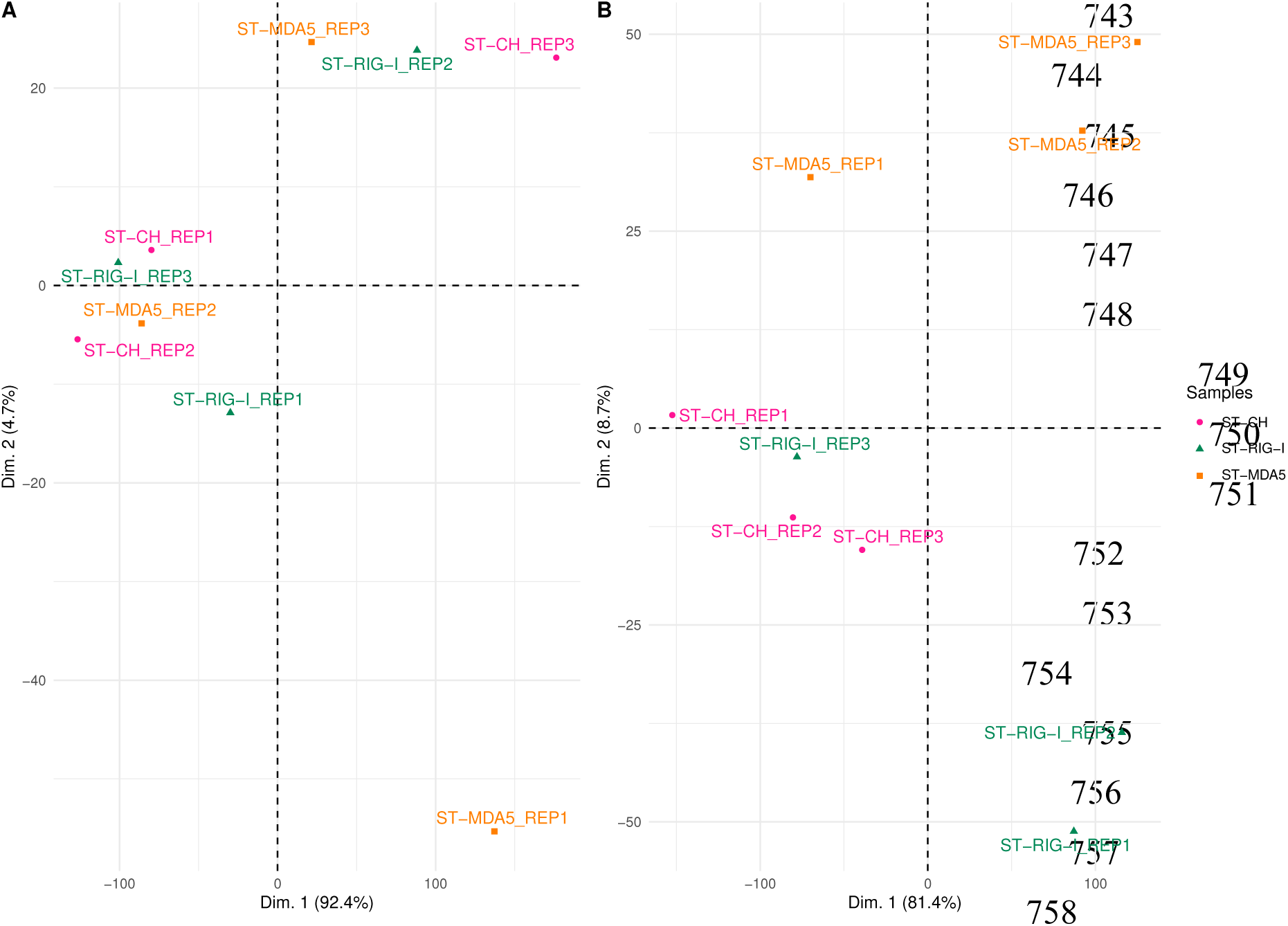
First two dimensions of principal component analysis (PCA) from normalized coverage for THA(A) and SAD (B) output samples. No group (or replicate) effect is detectable for THA. For SAD, differences between MDA5 and RIG-I against cherry, on the first dimension which explain 80 % of the variability.

**Figure S3:**
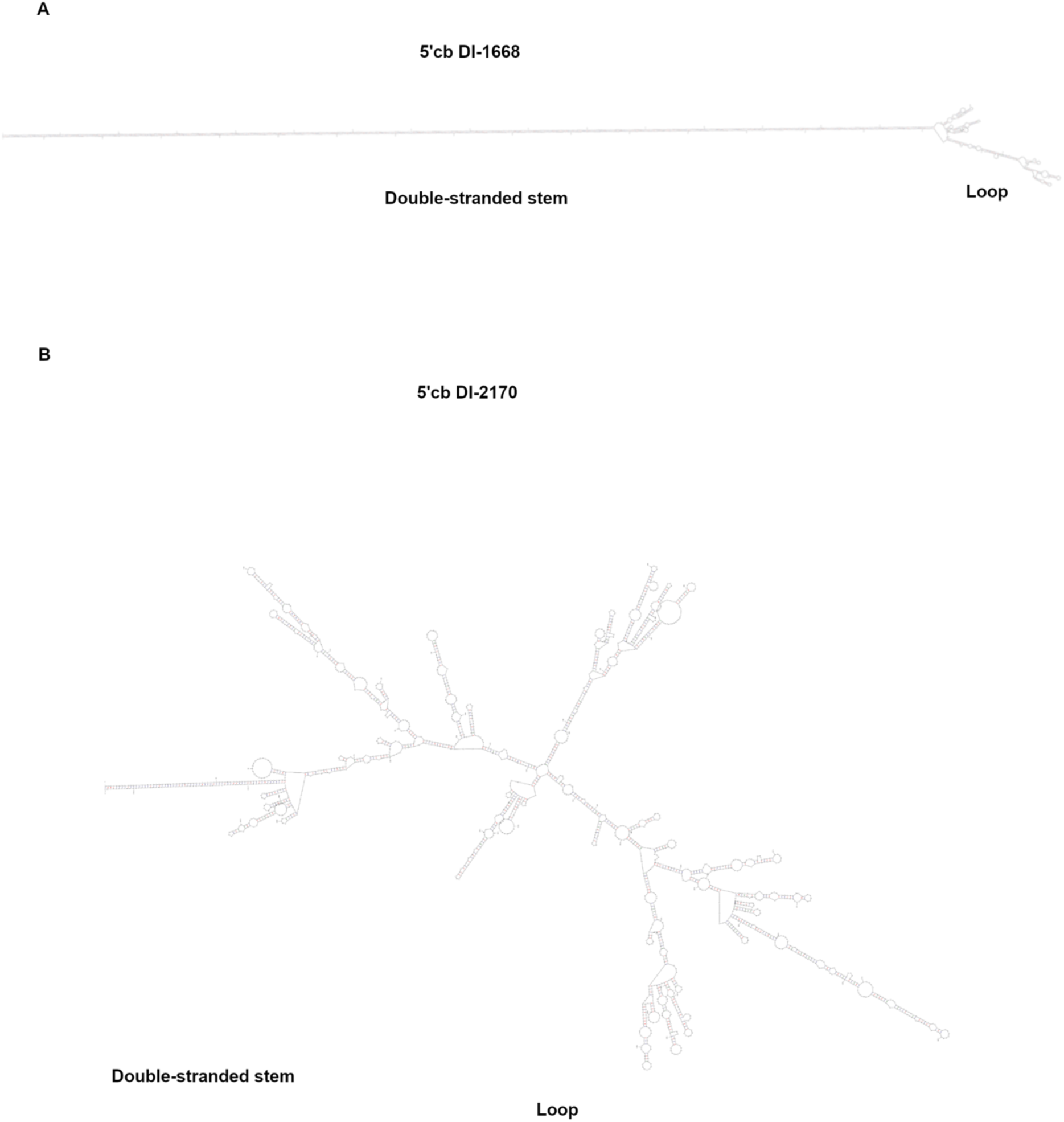
mFold secondary structure prediction of copy-back DI RNAs.

**Figure S4:**
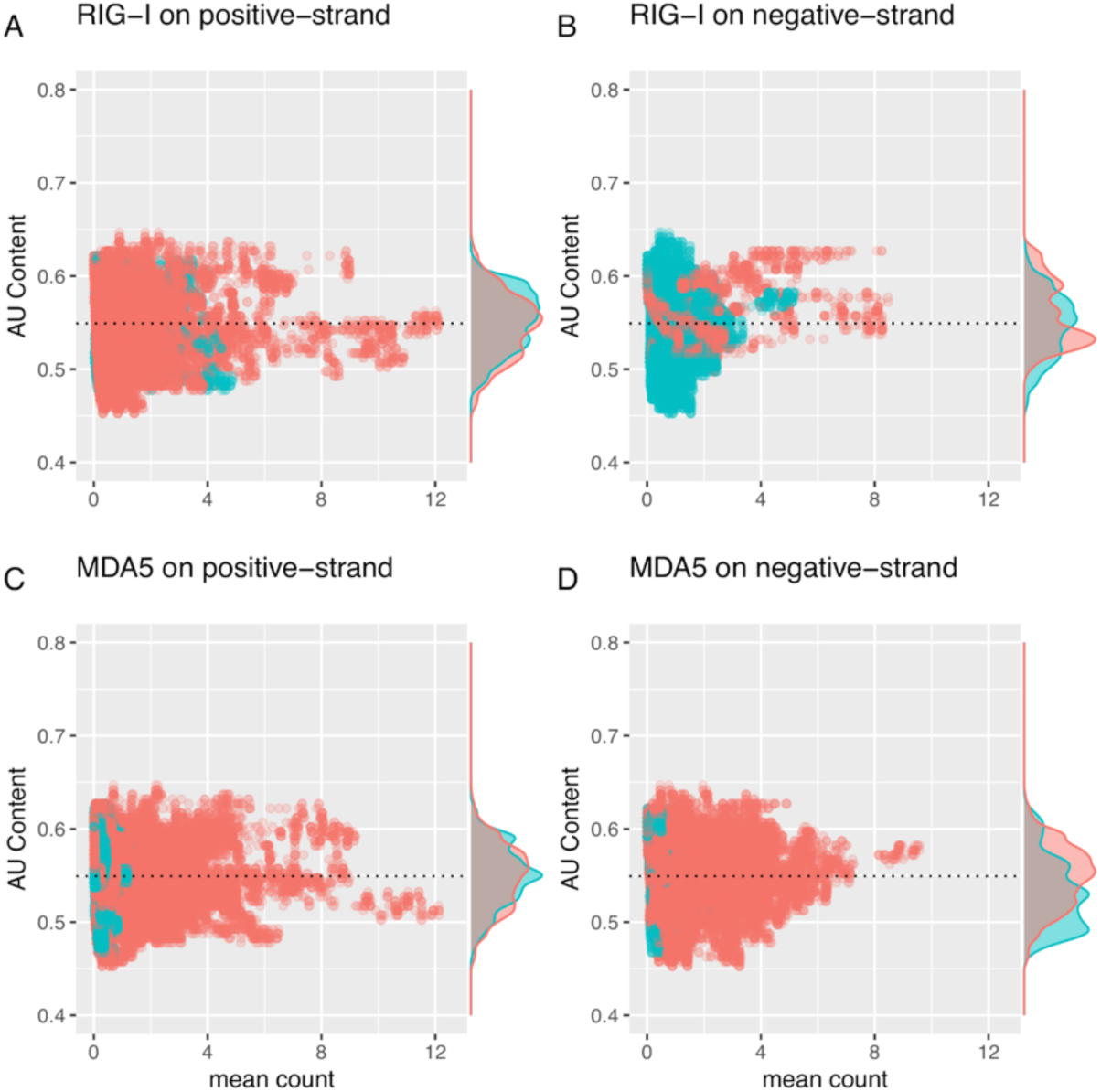
*In silico* analysis of RLR-specific SAD ligands. AU content of specific RLR RNAs partners (A, B) for RIG-I and (C, D) for MDA5. Relation between AU content and mean count for each position. Significant binding positions (Fold change > 2) are represented in orange and non-significant are colored in blue. Marginal distribution of AU content is added as density plot. Dashed line represents the AU content of SAD genome (%AU=0.55).

